# *Shigella* generates distinct IAM subpopulations during epithelial cell invasion to promote efficient intracellular niche formation

**DOI:** 10.1101/2023.05.22.540987

**Authors:** Lisa Sanchez, Michael G. Connor, Mélanie Hamon, Camila Valenzuela, Jost Enninga

## Abstract

The facultative intracellular pathogen *Shigella flexneri* invades non-phagocytic epithelial gut cells. Through a syringe-like apparatus called type 3 secretion system, it injects effector proteins into the host cell triggering actin rearrangements leading to its uptake within a tight vacuole, termed the bacterial-containing vacuole (BCV). Simultaneously, *Shigella* induces the formation of large vesicles around the entry site, which we refer to as infection-associated macropinosomes (IAMs). After entry, *Shigella* ruptures the BCV and escapes into the host cytosol by disassembling the BCV remnants. Previously, IAM formation has been shown to be required for efficient BCV escape, but the molecular events associated with BCV disassembly have remained unclear. To identify host components required for BCV disassembly, we performed a microscopy-based screen to monitor the recruitment of BAR domain-containing proteins, which are a family of host proteins involved in membrane shaping and sensing (e.g. endocytosis and recycling) during *Shigella* epithelial cell invasion. We identified endosomal recycling BAR protein Sorting Nexin-8 (SNX8) localized to IAMs in a PI(3)P-dependent manner before BCV disassembly. At least two distinct IAM subpopulations around the BCV were found, either being recycled back to cellular compartments such as the plasma membrane or transitioning to become RAB11A positive “contact-IAMs” involved in promoting BCV rupture. The IAM subpopulation duality was marked by the exclusive recruitment of either SNX8 or RAB11A. Finally, hindering PI(3)P production at the IAMs led to an inhibition of SNX8 recruitment at these compartments and delayed both, the step of BCV rupture time and successive BCV disassembly. Overall, our work sheds light on how *Shigella* establishes its intracellular niche through the subversion of a specific set of IAMs.

## Introduction

The entero-invasive bacterial pathogen *Shigella flexneri* (hereafter referred to as *Shigella*) is the causative agent of bacillary dysentery which affects an estimated 80 to 165 million individuals annually (CDC, 2019) representing a major health threat (Mahbubur et al. 2007, Kim et al. 2008, Puzari, Sharma and Chetia, 2018). To invade non-phagocytic epithelial gut cells *Shigella* uses a syringe-like apparatus termed the type 3 secretion system (T3SS) to reprogram the host actin cytoskeleton around the entry site (Schroeder and Hilbi, 2008, Valencia-Gallardo et al. 2014). This leads to the formation of positively shaped membranes ruffles whose collapse prompts the formation of concave membrane compartments. These enable (i) the internalization of the bacterium within a tight phagosome-like vacuole (Weiner et al. 2016) referred to as the bacteria-containing vacuole (BCV) and simultaneously (ii) the formation of heterogeneous vesicles in size with a similar morphology to macropinosomes (Cossart and Sansonetti, 2004, Weiner et al. 2016), termed infection-associated macropinosomes (IAMs). *Shigella* then triggers BCV rupture and disassembly prompting its access to the host cell cytosol where it replicates and spreads by forming an actin comet tail propelling it to adjacent cells (Cossart and Sansonetti, 2004, Kühn et al. 2020, Chang et al. 2020).

Although the contribution of host molecular pathways subverted in the invasion and niche establishment of intravacuolar bacterial pathogens is well-defined, it remains less clear for cytosol-residing bacteria (López-Montero and Enninga, 2016, Mellouk and Enninga, 2016). While BCV rupture has been suggested to be T3SS-mediated through the translocator proteins IpaB and IpaC (High et al. 1982, Blocker et al., 1999, Du et al. 2012), we previously demonstrated an impact of the endosomal recycling small GTPase RAB11A recruited to IAMs in promoting BCV rupture (Mellouk et al. 2014, Weiner et al. 2016). We furthermore reported that BCV damage is followed by membrane remnant disassembly, a process indispensable for *Shigella* cytosolic niche establishment (Kühn et al. 2020, Chang et al. 2020). Membrane disassembly requires RAB8A and RAB11A to be recruited to IAMs and brought in proximity of the BCV by the exocyst complex (Chang et al. 2020). Combined, these findings demonstrate a need to comprehensively analyze the contribution of host trafficking factors in the context of *Shigella* invasion. Moreover, they depict IAMs not as bystander compartments but rather as main actors in the process of *Shigella* intracellular niche formation highlighting the need to revisit and clarify the *Shigella* infection process.

Membrane remodeling is actively driven by a combination of local changes in membrane lipid composition and protein-generated membrane reshaping. This process drives the formation of a positive or a negative curvature necessary for several cellular processes from filopodia formation to endocytosis (McMahon and Gallop, 2005, Jarsch et al. 2016, Simunovic et al. 2019). Several bacterial invasion processes induce extensive host membrane reshaping by curving the membrane surrounding the entering bacteria prior to subsequent curvature collapse, as exemplified by membrane ruffling during *Shigella* entry (Swanson, 2008, Cossart and Roy, 2010, Ribet and Cossart, 2015, Buckley and King, 2017).

Moreover, the formation of phagosomes and macropinosomes have also been shown to require extensive membrane reshaping (Swanson, 2008) for vesicle formation, scission and stability which all require changes in lipid composition as well as protein intervention (Swanson, 2008, Swanson, 2014).

BAR (Bin/Amphiphysin/Rvs) domain-containing proteins have been described as membrane-binding proteins specializing in membrane reshaping and curvature sensing (McMahon and Gallop, 2005, Allison Suarez et al. 2014, Simunovic et al. 2015, Simunovic et al. 2019). Characterized by the presence of a membrane-binding BAR domain, these proteins act in diverse cellular processes as signaling platforms (*e.g.* filopodia formation) and scaffolds (*e.g.* endosomal recycling) (Peter et al. 2004, Simunovic et al. 2015, Simunovic et al. 2019). Due to this, BAR domain proteins have been reportedly hijacked by invasive bacterial pathogens to facilitate their entry into the host cell. *Enterohemorrhagic E. coli* was reported to reprogram negative curvature-inducing and actin remodeling factor IRSp53 to form an actin structure called a pedestal (Weiss et al. 2009, Yi and Goldberg, 2009).

Similarly, endosomal recycling BAR protein SNX1 is recruited to the *Salmonella*-containing vacuole through the bacterial effector SopB to form tubular structures forming the *Salmonella* replicative niche (Bujny et al. 2008, Stévenin et al. 2019). In the case of *Shigella* TOCA-1 has been implicated in actin rearrangements for the formation of an actin cocoon prior to BCV rupture, and it is present at actin tails (Leung et al. 2008, Baxt and Goldberg, 2014, Kühn et al. 2020).

Here, we exploited a high-content multidimensional time-resolved fluorescence microscopy assay (Sanchez et al. 2021) to screen a BAR protein library and comprehensively analyze their involvement during the successive *Shigella* invasion steps. We identified the sorting nexin-BAR family member SNX8 to be strongly present at a subset of IAMs before BCV damage and remain there until membrane remnant disassembly. The characterization of the SNX8-positive subset showed a distinct maturation of IAMs from “canonical” macropinosomes. Furthermore, we found this IAM subset to promote efficient BCV egress. This diversity in IAMs reveals the *Shigella* invasion process as a complex sequence of events with the bacteria hijacking multiple pathways of the endocytic and recycling pathways for bacterial survival and cytosolic access.

## Materials and Methods

### Bacterial strains and culture

In this study we used the *Shigella flexneri* strain M90T (Sansonetti et al. 1982) expressing the uropathogenic *E. coli* adhesin AfaI. Prior to infection experiments, *Shigella* strains were grown overnight at 37°C from bacterial colonies grown on Trypticase Casein Soy Broth agar plates supplemented with ampicillin at 50µg/mL and 0.01% Congo Red. Bacterial cultures were prepared by inoculating 3 colonies in TCSB media supplemented with ampicillin 50µg/mL and incubated overnight at 37°C, 220 rpm.

### Cell lines and culture conditions

HeLa cervical adenocarcinoma CCL2 clone from the American Type Culture Collection (ATCC) and CaCo-2 TC7 cells (ATCC) were cultured in Dulbecco’s Modified Eagle Medium (DMEM High glucose with GlutaMAX^TM^ and pyruvate, Gibco, #31966-021) supplemented with 10% heat-inactivated Fetal Bovine Serum (Sigma Aldrich) and incubated at 37°C, 5% CO2.

### Plasmids, cloning and cell line generation

The full list of plasmids used in this study is listed in **Supplementary Table 1**. The entire EGFP-tagged BAR domain plasmid library was a kind gift from Emmanuel Boucrot and is referenced in Chan Wah Hak et al. 2018. pDEST-SNX8-mApple was cloned by restriction enzyme digestion. Generation of stable HeLa cell lines was performed using the Sleeping Beauty System (Kowarz et al. 2015). We generated cell lines expressing fluorescent SNX8 and the plasma membrane marker LactC2 using pSBbi-Neo-SNX8-eGFP and pSBbi-Neo- LactC2-GFP. Both were cloned by *in vivo* assembly using SLIC (Jeong et al. 2012). JetPRIME (Polyplus, #101000027) was used to transfect plasmids into low passage HeLa cells with selection being performed using G418 at 800ng/mL (Euromedex, #EU0601) for 7 days. The cells were then collected, and serial dilution was performed in a 96 well plate with maintenance of the selection pressure. The selected cells were then amplified and sorted based on their fluorescence level using a BD FACSAria^TM^ III Cell Sorter. Live cells were gated in FSC-A/SSC-A and doublets were removed using the parameters FSC-A and FSC-H. Three populations, according to the GFP or EGFP intensities, were sorted as low-, medium- and high-expressing GFP/EGFP cells.

### Cell seeding and Transient transfections

Cells were seeded in either black 96-well plates (Greiner Bio-One, #655090) or in a 35mm glass-bottom dish (ibiDI, #81158) containing a 4-chamber silicone insert (ibiDI, #80409). For both supports, HeLa cells were seeded at a density of 8,000 cells per well, whereas CaCo-2 TC7 cells were seeded at a density of 5,000 cells per well. Transient transfections were performed the following day by lipofection using FuGENE^TM^ HD (Promega, #E2311) as instructed by the manufacturer. Briefly, transfection complexes were prepared by diluting 2 µg of plasmid (or for co-transfections, 1µg of each plasmid) in 100µL OptiMEM (Gibco, #31985062), mixed with 4 µL of FuGENE^TM^ HD and incubated for 10 min at room temperature. Afterwards, 5 µL of transfection mixture was added per well containing the cells seeded the previous day, the cells were then incubated 48 hours at 37°C, 5% CO2 prior to infection and imaging.

### Infection protocol

Infections were carried out as previously described (Chang et al., 2020). Bacterial inoculum were prepared by diluting the overnight culture to 1:100 in 8 mL of fresh TCSB media supplemented with ampicillin 50µg/mL and incubation for 2 hours at 37°C, 220 rpm. Once the subculture OD600nm reached 0.4-0.6, 1mL of the subculture was spun 1 min at 6000*g* and the bacteria pellet was washed twice with warm EM buffer (120 mM NaCl, 7 mM KCl, 1.8 mM CaCl2, 0.8 mM MgCl2, 5 mM glucose, 25 mM HEPES, pH 7.3). An inoculum was prepared by diluting the bacterial suspension in EM buffer to reach a multiplicity of infection (MOI) of 20 bacteria/cell. For inhibitor experiments, wortmannin (Sigma Aldrich, #W1628-1MG) and SAR405 (Selleckchem, #S7682) were diluted into the inoculum to 3.3µm and 3µM respectively from a DMSO stock solution. Prior to infection, the cells were washed three times with EM Buffer, and 50µL of EM buffer were left in the well. For time lapse imaging experiments, the infection was started by adding 40 µL of inoculum per well in a 37°C heated microscopy chamber and image acquisition was started. For endpoint experiments through cell fixation, no medium was left in the well and 30 µL of inoculum plus 0.5mg/mL final dextran 10 000 MW Alexa-647 (Invitrogen^TM^, #D22914) were added to the well, after which the samples were incubated at 20°C for 10 min to enable the bacteria to reach the cells. Infection was triggered by incubation at 37°C for 30min. Afterwards, samples were fixed using 4% paraformaldehyde (ThermoScientific, #043368.9M) for 10 min at room temperature and washed 3 times with PBS. DNA was stained using Hoechst (Invitrogen, #1681305) and actin was stained by rhodamine-phalloidin (Invitrogen, #R415) for 30 min prior to microscopy acquisition.

### Immunofluorescence

Following the fixation procedure, the cells were permeabilized using saponin diluted to 0.025% in PBS for 20 min at room temperature. Next, 3 PBS washes were performed and blocking was done with 2% Bovine Serum Albumin (Sigma Aldrich,#A7906-100G) and 5% goat serum (Sigma Aldrich, #G9023) in PBS for 1 hour at room temperature. Afterwards, rabbit anti-SNX8 primary antibody (Sigma Aldrich, #HPA057296) diluted at 1:250 in blocking solution was incubated 1h at room temperature followed by a 45min incubation of a goat anti-rabbit Alexa-488 secondary antibody (Invitrogen, #A11034) diluted to 1:500 together with Hoechst and rhodamine-phalloidin. Finally, cells were washed 3 times with PBS and left in PBS until imaging.

### Time lapse microscopy

The time-resolved BAR domain protein screen experiments were performed in a Nikon Ti-E inverted microscope equipped with a Perfect Focus System (TI-ND6-PFS Perfect Focus Unit) using a 40X/ 0.75 NA air objective. High spatio-temporal resolution time-lapses were acquired on DeltaVision Elite (Leica) using a 60X/1.42 NA oil objective with a 0.35µm z-step and images were deconvolved using an integrated deconvolution software. Imaging of fixed experiments were acquired on a Nikon Ti-E inverted microscope equipped with a Perfect Focus System and a Yokogawa confocal spinning disk unit (CSU-W1) using a 60X/1.2 NA water objective. In this case, an automatic pipeline with autofocus using brightfield and Hoechst signal were used to define the focal plane of randomly generated positions. Images were acquired at a step-size of 0.5µm in the z-plane.

### Image Processing and quantification

Images were processed using Fiji (https://imagej.net/software/fiji/, version 2.1.0/1.53c). For the BAR domain screen, BAR protein TOCA-1 and EGFP were used as positive and negative controls respectively (Leung et al. 2008, Kühn et al. 2020). Positive hits were counted as showing enrichment of the “candidate” proteins to the infection site by comparison to the EGFP control (see **Figure 1B**). For the quantification of SNX8 positive IAMs, the SNX8 cytosolic fluorescence background was subtracted from the image. Cell Profiler (www.cellprofiler.org, version 4.2.1) was used to count SNX8 positive IAMs. Briefly, following removal of the cytosolic fraction of SNX8 in the SNX8 channel using Fiji, IAMs were detected using the dextran signal and the dextran-IAMs were used as a mask on the SNX8 signal. The software parameters were set to detect the upper quartile fluorescence intensity of the SNX8 to the IAMs and this was related to the detected bacteria and actin foci.

**Figure 1.**
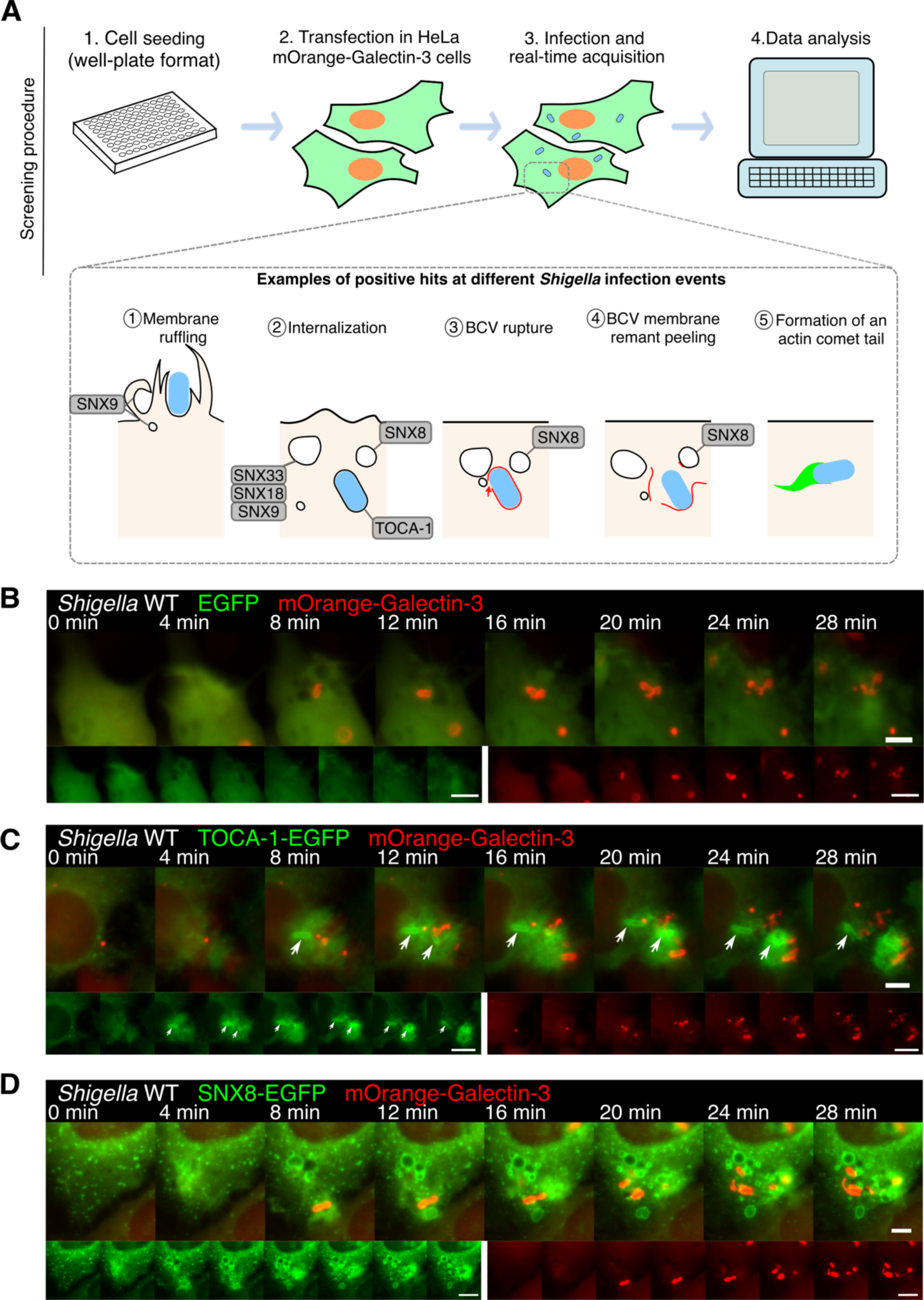
Time-resolved high-content screen experimental procedure and examples of positive hits of host proteins recruited to the bacterial infection foci. **A.** Schematic illustration of the time-resolved screen workflow with example of positive hits and their observed localization during *Shigella* infection events. HeLa mOrange-Galectin-3 cells were seeded, transfected with the proteins-of-interest and samples were infected with *Shigella* during microscopy acquisition. Examples of observed hit recruitment and their localization to the infection site are represented in the lower portion the figure. **B. C. D.** Microscopy images of external controls EGFP and TOCA-1 as well as an example of positively identified hit: the SNX-BAR family SNX8 protein. In all figures, in red is shown the mOrange-Galectin-3 signal marking *Shigella*-BCV rupture and the BCV-membrane remnants, and in green the BAR-domain containing proteins included in the screen. White arrows show the recruitment of TOCA-1-EGFP to un-ruptured BCVs. Scale bars are 5µm and 10µm.

### Statistical analysis

Statistical analysis was performed using GraphPad Prism version 9.0.0 for MacOS, GraphPad Software, San Diego, California USA, www.graphpad.com.

## Results

### Identification of BAR domain-containing host factors enriched at *Shigella* invasion foci

With the aim of identifying new molecular pathways involved in the early steps of *Shigella* invasion of epithelial cells, we carried out a high-content time-resolved microscopy screen using a library of 66 EGFP-tagged full length BAR domain-containing proteins (see **Supplementary Table 1** for full list). In parallel, we also followed the signals of fluorescently tagged Galectin-3, a cytosolic reporter that binds to the inner leaflet of the BCV membrane at the precise moment of vacuole rupture (Paz et al. 2010). In brief, a 2-minute-interval time-course of mOrange-Galectin-3 stably expressing HeLa cells transiently transfected with each plasmid from the BAR protein library and challenged with wildtype (WT) *Shigella* were recorded (**Figure 1A**). This set-up enabled to track the BAR domain protein being recruited to the infection focus and simultaneously monitoring of 3 distinct *Shigella* invasion events: a) *Shigella*-triggered membrane ruffling, b) the step of BCV rupture, and c) the events of BCV membrane disassembly upon rupture. Candidate targets were scored against a positive recruitment control TOCA-1 (Leung et al. 2008, Kühn et al. 2020) and EGFP negative recruitment control. (**Figure 1B**).

After manual inspection of the data based on the behavior of the external controls, 13 BAR domain-containing proteins were identified as positive hits (see **Supplementary Table 2**), whereas 10 were determined to be inconclusive due to poor transfection efficiency or low fluorescence signal and were removed from further analysis. Our results revealed specific subtypes of BAR domain-containing proteins localizing to different bacterial compartments. Among the hits were several members of the SNX-BAR family of proteins and actin nucleating factors such as Oligophrenin and PACSINs 1-3. We observed distinct recruitment patterns for these hit proteins during the infection. While PACSIN3 recruitment was observed early during membrane ruffling, Oligophrenin was found to localize at the BCV prior to BCV rupture (**Supplementary Figure 1**). Interestingly, multiple members of the SNX-BAR family localized to IAMs early in their formation and/or their enrichment occurred throughout *Shigella* BCV egress (**Figure 1** and **Supplementary Figure 1**). Among them, a significant enrichment of the early endosome sorting protein Sorting Nexin 8 (SNX8) to IAMs (**Figure 1D**). This recruitment was observed early on during the infection, and the protein remained persistently localized at the IAMs throughout the steps of *Shigella* BCV disassembly. Giving the role of IAMs in BCV rupture and unpeeling, the persistent recruitment of SNX8 suggest it could be involved in several steps leading to cytosolic access.

### SNX8 localizes to IAMs and the *Shigella*-BCV prior to BCV rupture and disassembly

SNX8 has been shown to localize to early endosome and suggested to recycle endocytic cargo to the Trans-Golgi network (Dyve et al. 2009). Although the function of human SNX8 remains elusive, its yeast homologue MVP1 has been shown to function in retromer-independent recycling (Suzuki et al. 2021). SNX8 has also been linked to several pathologies involving endosomal recycling defects such as Alzheimer’s (Xie et al. 2019, Vanzo et al. 2014). With our identification of SNX8 localization and retention to IAMs we sought to define its functional contribution to *Shigella* entry.

Following the identification of SNX8 as a hit in our screen, we proceeded to comprehensively characterize SNX8 recruitment to IAMs. To rule out SNX8 recruitment to the BCV or IAMs due to ectopic expression artifacts we determined the localization of endogenous SNX8 by immunodetection using an anti-SNX8 antibody. Simultaneously, we monitored all of the plasma membrane-derived IAMs employing the previously described phosphatidylserine-specific biosensor LactC2 (Yeung et al. 2008, Vecchio and Stahelin 2018). Confocal microscopy analysis confirmed the presence of endogenous SNX8 to LactC2- marked vesicles in proximity to *Shigella* (**Figure 2A**). We then assessed SNX8 behavior in relation to the ruptured BCV membrane. Temporal analysis of images from higher spatial resolution live-cell imaging (see methods for details) of HeLa and CaCo-2 cells transiently co-expressing SNX8-EGFP and mOrange-Galectin-3 showed the recruitment of SNX8 to *Shigella*- IAMs occurred prior to BCV rupture and remained at the IAMs until full cytosolic release of *Shigella* after BCV disassembly (**Figures 2B** and **2C**). Our results show SNX8 recruitment to the *Shigella* entry foci roughly 6 min prior to BCV rupture and throughout BCV disassembly, which we previously showed occurs 20min post-infection Chang et al. 2020) (**Figure 2D**).

**Figure 2.**
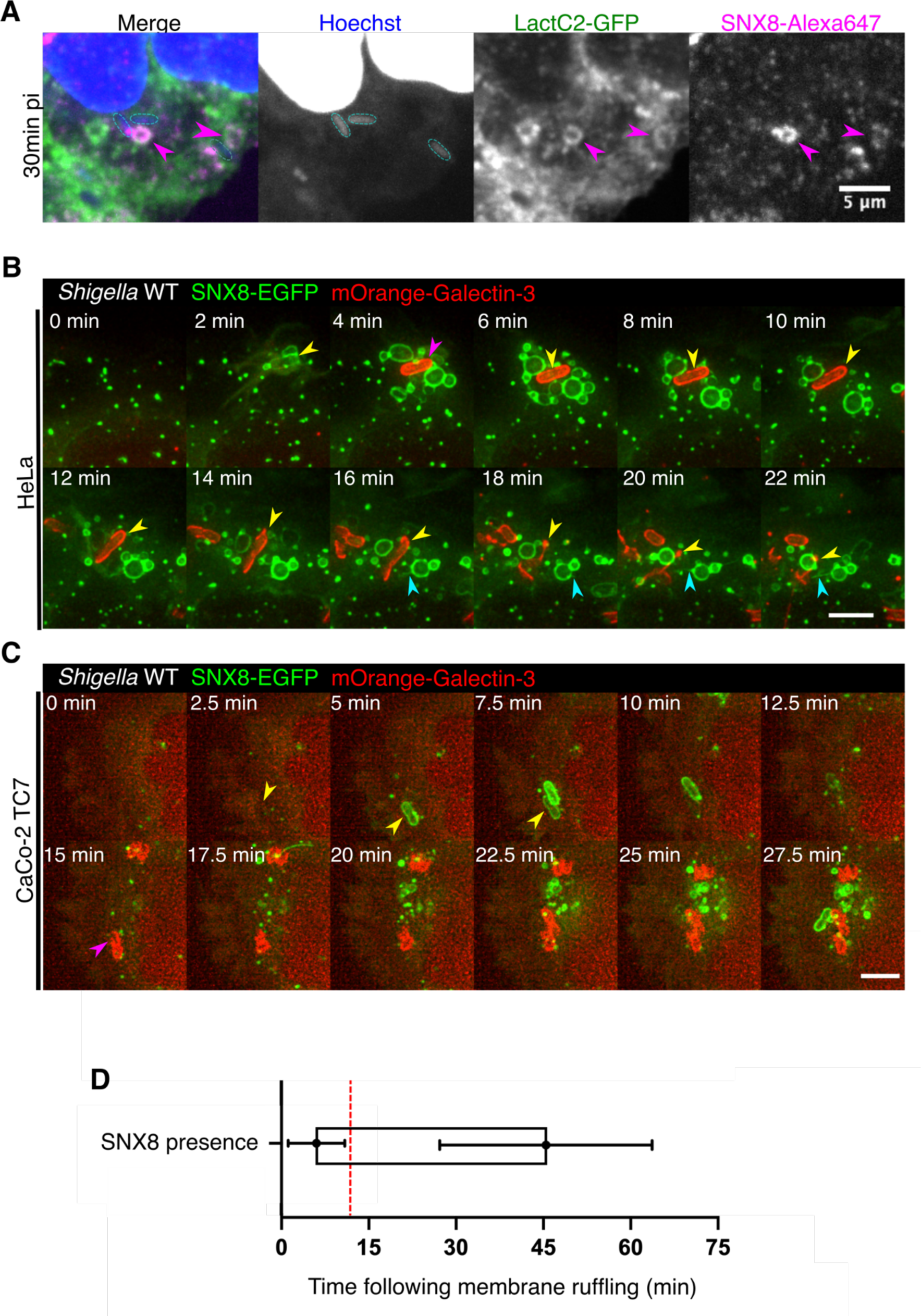
SNX8 is recruited to IAMs and BCV prior to *Shigella*-BCV egress. **A.** Fixed confocal microscopy of LactC2-GFP stably expressing HeLa cells (in green) infected with *Shigella* for 30 min and stained for endogenous SNX8 (in magenta). The bacteria and cell nucleus were marked with Hoechst (in blue). Magenta arrowheads show individual IAMs recruiting endogenous SNX8 and bacteria are outlined in cyan. Scale bar: 5µm. **B, C.** Time-lapse microscopy images of HeLa cells (B) and CaCo-2 cells (C) transfected with mOrange-Galectin-3 (in red) and SNX8-EGFP (in green) infected with *Shigella* WT. Yellow arrowheads point to an entering bacterium, magenta arrowheads indicates the moment of BCV rupture. SNX8-rich tubules emanating from SNX8-IAMs are shown by cyan arrows. The infection start (t=0) was defined by the first apparition of membrane ruffles. Scale bar: 3µm. **D.** Quantitative analysis of SNX8-EGFP temporal recruitment sequence to the infection focus. SNX8 presence is shown as a box with the average start of the recruitment at 6min and dissipation at 44.8min. Infection foci from two biological replicas were analyzed (n>60). Bar shows the average start and end of SNX8 presence at the BCV +/- standard deviation. The red line represents the average BCV rupture time point (11.9min).

Moreover, SNX8 time-lapses showed SNX8 positive tubules emanating from IAMs (**Figure 2B**), as previously described for SNX-BARs by Van Weering et al (2010). We also observed the BCV to be transiently enriched in SNX8 prior to BCV rupture (**Figure 2B**). Altogether, these results temporally map SNX8 recruitment to IAMs at early steps of BCV rupture and/or disassembly.

### SNX8 recruitment is partially-driven by *Shigella* bacterial effectors IpgD and IcsB

Our data demonstrated SNX8 is recruited to IAMs prior to BCV rupture/disassembly, a process known to require T3SS secreted *Shigella* effectors. Thus, we investigated if SNX8 localization was also effector dependent. Given the rapid recruitment of SNX8 to IAMs we investigated the impact of the first set of secreted bacterial effector proteins IpgD and IcsB. The *Shigella* phosphatase, IpgD, is involved in the recruitment of PI(3)P to IAMs and mutants have been reported to decrease IAM numbers with delay of BCV rupture (Mellouk et al. 2014, Weiner et al. 2016). The effector IcsB, an N-fatty acylase, is known to modify multiple cellular host factors (Liu et al. 2018) and participates in actin cocoon formation (Kühn et al, 2020).

In order to monitor SNX8 dynamics or recruitment a 2-minute interval microscopy time-course was performed using SNX8-EGFP and mOrange-Galectin-3 co-transfected HeLa cells infected with either wildtype *Shigella* or deletion mutants *of* Δ*icsB* and Δ*ipgD*. Analysis of these time lapse movies showed a notable decrease in the SNX8-IAMs surrounding the BCV upon invasion of the *Shigella* Δ*ipgD* or Δ*icsB* mutants (**Figures 3B** and **3C** respectively) compared to the *Shigella* WT infected cells (**Figure 3A**). Additionally, *Shigella* Δ*ipaJ* mutant infections were performed resulting in a similar phenotype as *Shigella* WT infections (**Supplementary Figure 2**). This demonstrates a specificity of the Δ*ipgD* and Δ*icsB* effectors for the recruitment of SNX8 to IAMs and indicates that recruitment of SNX8 is driven by *Shigella* through them.

**Figure 3.**
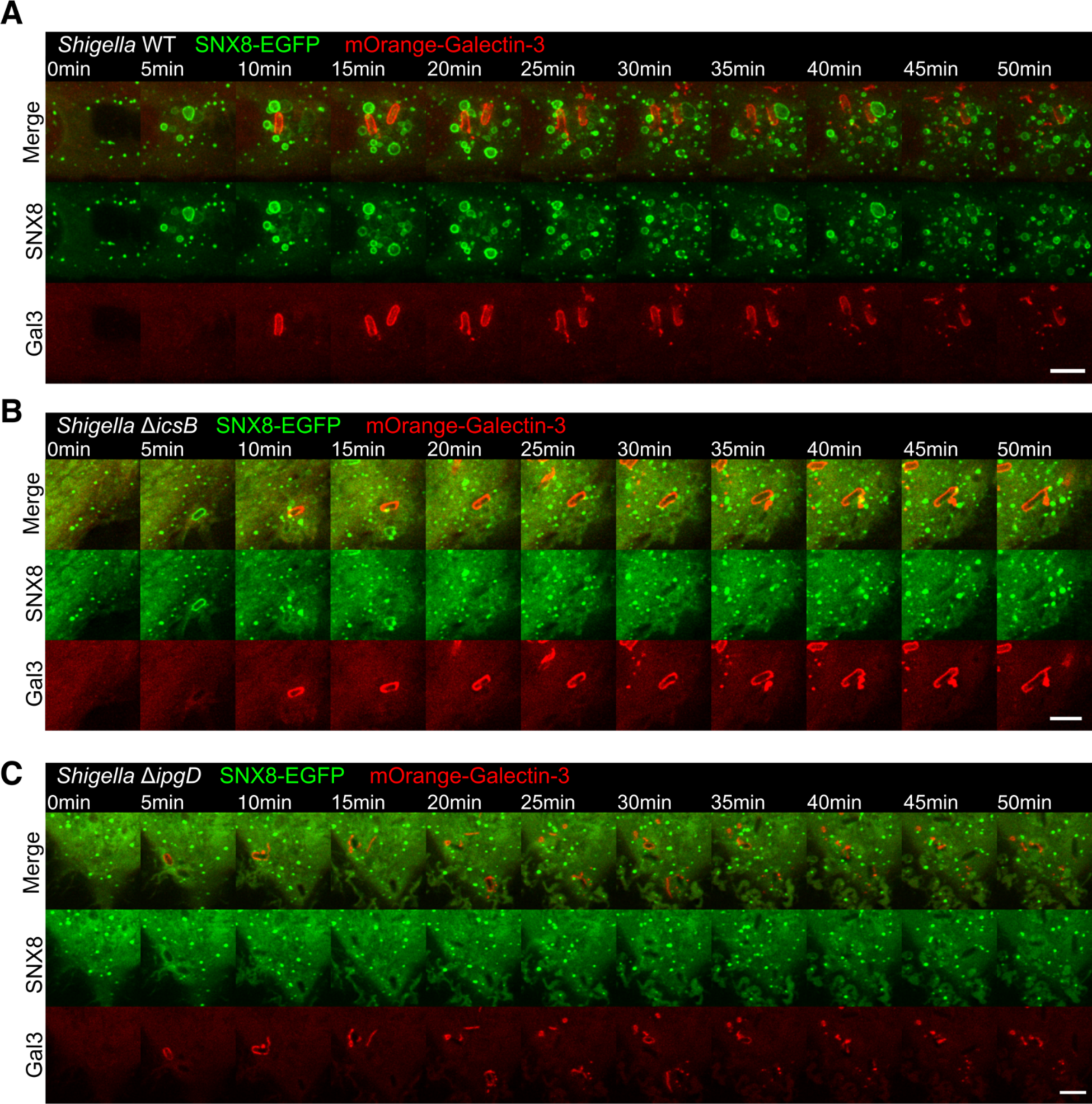
SNX8 recruitment to the *Shigella* infection site is impacted by the bacterial effectors IpgD and icsB. HeLa cells co-expressing SNX8-EGFP (in green) and mOrange-Galectin-3 (in red) were infected with either *Shigella* WT (**A**) or the mutants Δ*ipgD* (**B**) and Δ*icsB* (**C**). Galectin-3 signal indicates the moment of BCV rupture and labels the ruptured BCV membrane-remnants. The infection start (t=0) was defined by the first apparition of membrane ruffles. Scale bar is 5µm.

### SNX8 is recruited to a PI(3)P positive subpopulation of IAMs

We previously showed that IAMs produced during *Shigella* infections are positive for PI(3)P (Weiner et al. 2016). Interestingly, SNX8 contains a PX domain binding to PI(3)P (Van Weering et al. 2012). We speculated that SNX8 recruitment and PI(3)P presence to IAMs occurred concurrently in time. To address this we performed time-lapse microscopy at 1 min-intervals to capture the *Shigella* infection of transiently co-transfected HeLa cells with SNX8-EGFP and mCherry-2xFYVE- a PI(3)P binding probe construct (Stenmark et al. 1996). Indeed, our microscopy showed the localization of 2xFYVE to IAMs occurred simultaneously with EGFP-SNX8 enrichment. Moreover, SNX8 recruitment occurred to all 2xFYVE-labelled IAMs (**Figure 4A**). This data shows accumulation of SNX8 is exclusive to a subpopulation of PI(3)P-enriched IAMs.

**Figure 4.**
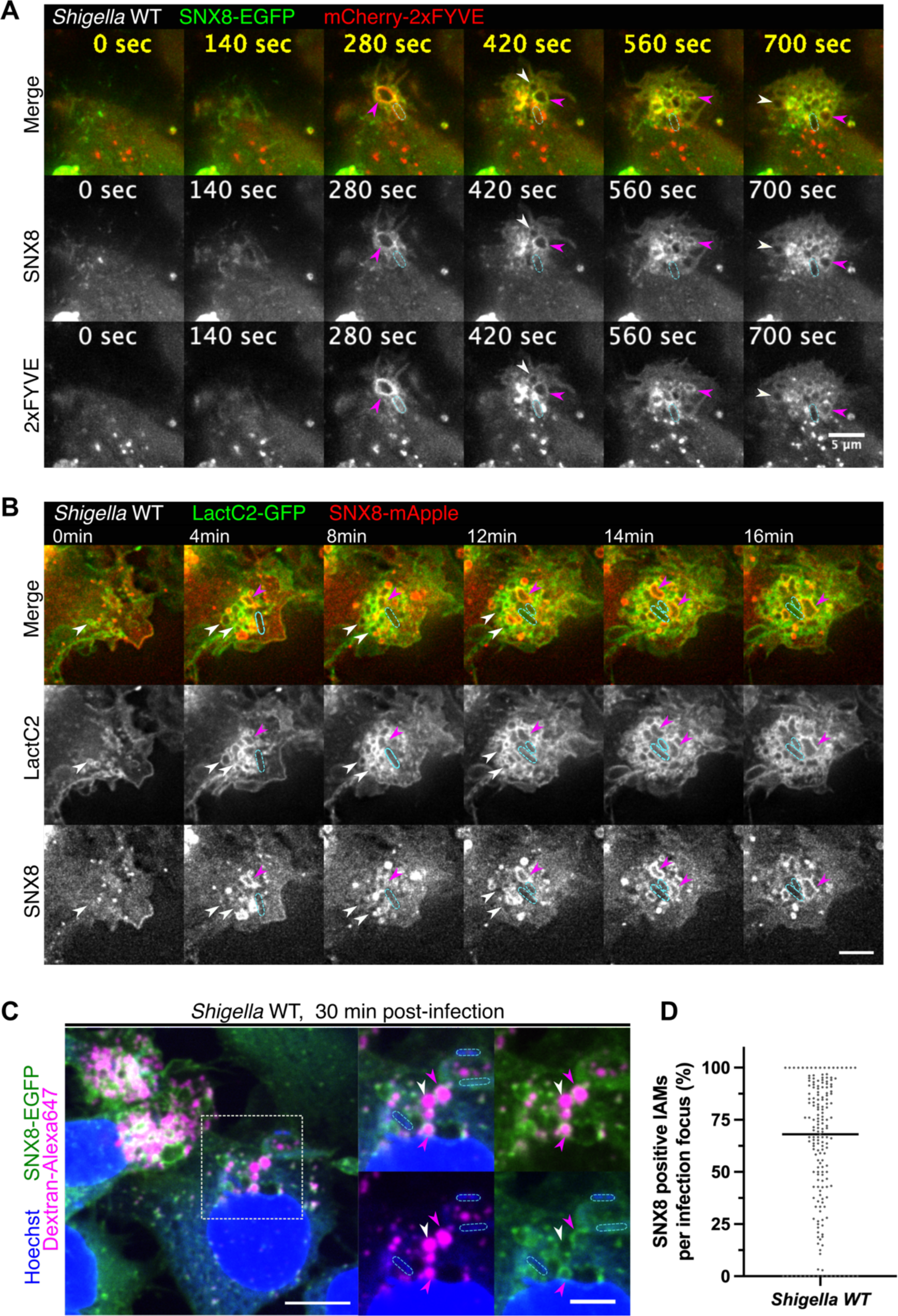
SNX8 is heterogeneously recruited to IAMs in a PI(3)P-dependent manner. Time-lapse microscopic analysis of SNX8 distribution at IAMs by transfecting SNX8-mApple in (**A**) cells co-transfected with 2xFYVE-EGFP or (**B**) LactC2-GFP-expressing HeLa cells. Entering bacteria are tracked with cyan outlines. Magenta and white arrowheads highlight SNX8 positive and negative IAMs respectively. The infection start (t=0) was defined by the first apparition of membrane ruffles. Scale bar is 5µm. **C**. Fixed confocal image of HeLa cells expressing SNX8-EGFP were infected with *Shigella* together with dextran-Alexa647 at 30min post-infection. SNX8 signal is in green and dextran-containing IAMs are in magenta. Magenta arrowheads show SNX8 positive-IAMs, white arrowheads show SNX8 negative IAMs and in cyan is outlined the bacteria. Inset show: a zoomed merged image with all channels (top left panel). The top right panel shows the SNX8-EGFP (in green) localizing to part of the IAMs formed (in magenta). Bacteria (outlined) together with IAMs are shown in the bottom left panel and an overview to the SNX8-EGFP recruitment on the bottom right panel. Scale bars are 10µm and 5µm. **D**. Quantification of dextran-labelled IAMs recruiting SNX8-EGFP following 30 min incubation with WT *Shigella*. The average percentage of IAMs recruiting SNX8 per infection foci is shown and the median is displayed (68% +/- 5,5). Automated analysis was performed on two biological replicates (n>90 infection foci per experiment).

How IAMs are involved in the intracellular trafficking of Shigella remains unclear, and it is unknown whether all IAMs are of the same composition (Mellouk et al. 2014, Weiner et al. 2016, Chang et al. 2020). To assess whether SNX8 localizes to all *Shigella*-IAMs, we first performed time-lapse microscopy experiments at higher spatial resolution using the LactC2 reporter to carefully monitor IAMs. In brief, 1-minute-interval time-course experiments were performed of SNX8-mApple was transiently transfected into LactC2-GFP stable HeLa cells infected with *Shigella.* Image analysis confirmed that all IAMs were labelled by LactC2- GFP throughout their lifetime (**Figure 4B**). Moreover, we also noted that SNX8 recruitment to IAMs begins shortly after IAM cup closure to a subset of the total IAMs formed (**Figure 4B**). To address this, we performed experiments using the fluorescent fluid phase marker dextran to label IAMs (Weiner et al. 2016) in SNX8-EGFP transiently transfected HeLa cells that were fixed after 30 min of bacterial challenge. These results further confirmed SNX8 to be enriched only to a fraction of the formed IAMs with just over half of them being SNX8 positive (61%, ±4.7%) (**Figures 4C** and **4D**). Together, these results reveal the co-existence of at least two subsets of IAMs of distinct composition within an infection focus.

### Characterization of SNX8 behavior during IAMs maturation

Previously, several small RAB GTPases were shown to be recruited during *Shigella*-IAM maturation with several of them playing a role in promoting the invasion steps (RAB8A, RAB11A), also marking the importance of IAM-recruited factors (Mellouk et al. 2014, Weiner et al. 2016, Chang et al. 2020). Given the recruitment of SNX8 to IAMs at early stages of infection, we postulated there was a spatial-temporal relationship with the recruitment of Rab GTPases to the IAMs. To address this, we co-transfected SNA8 and RAB5A, RAB7A, RAB8A or RAB11A and performed time-lapse microscopy at 35 sec-interval to temporally resolve the precise recruitment of these different factors.

Canonical macropinosome maturation has previously been reported to involve the early marker RAB5A and late marker RAB7A (Egami et al. 2014, Buckley and King, 2017). Our data showed recruitment of RAB5A to IAMs early on during the *Shigella* invasion process as previously reported by Mellouk et al (2014) (**Figure 5A**). Microscopy time-lapse images showed SNX8 recruitment occurs simultaneously with RAB5A recruitment to IAMs (**Figure 5A** and **5B**). Furthermore, temporal analysis of SNX8 and RAB5A fluorescence intensity to individual IAMs further emphasized this observation (see **Figure 5B**). Microscopic analysis also revealed SNX8 recruitment to IAMs occurred prior to RAB7A accumulation and decreased with RAB7A presence (**Figure 5C** and **5D**). Moreover, nearly all SNX8-IAMs acquired RAB7A and we observed formation of SNX8-negative RAB7A-positive tubules suggesting distinct recycling pathways of RAB7 and SNX8. These results show SNX8 recruitment to occur in “canonical macropinosome”-like maturing IAMs as soon as the early stage of IAMs maturation and it remains present until later stages of IAM maturation.

**Figure 5.**
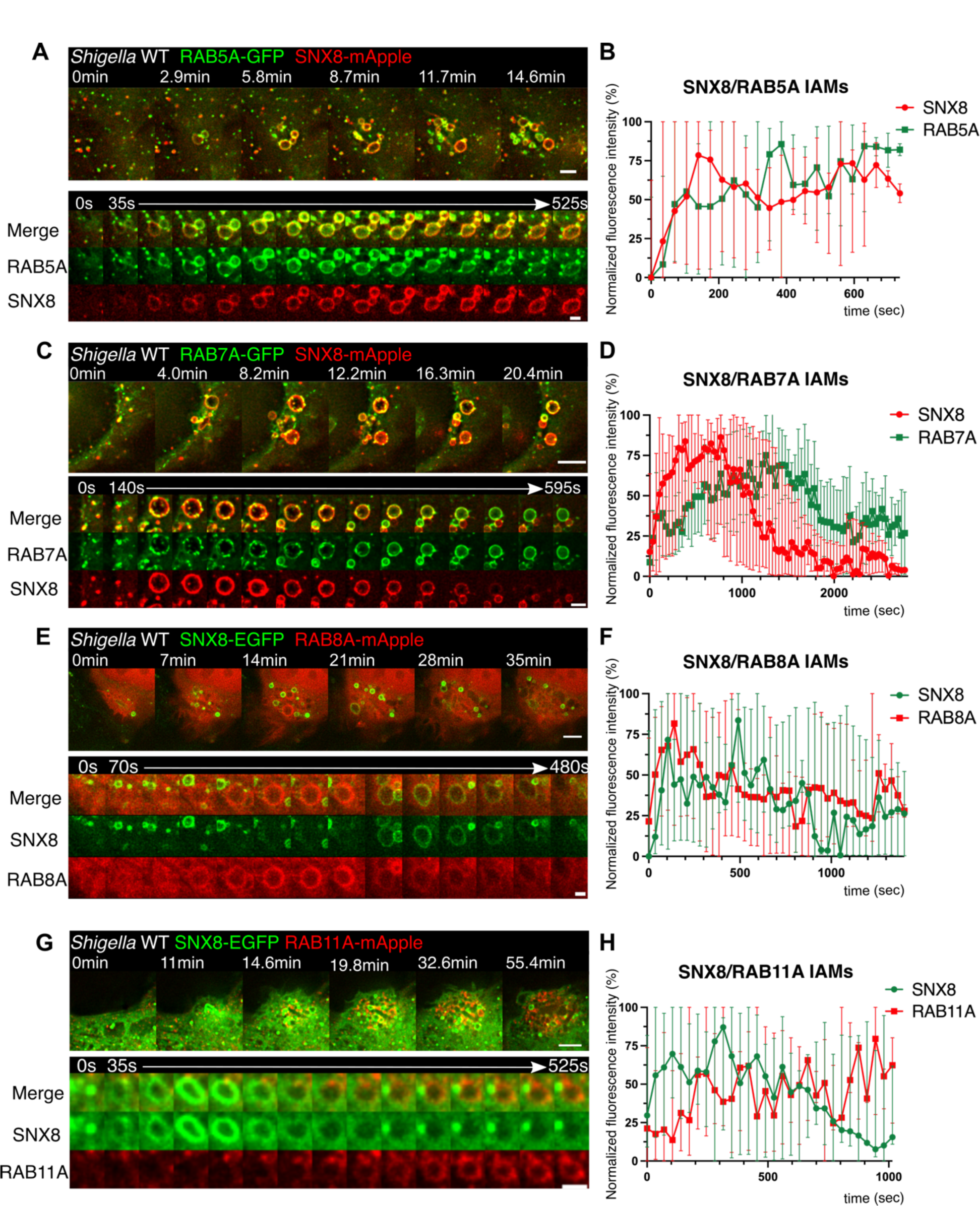
Characterization of the maturation of SNX8-IAMs to RAB GTPases. Live cell 35sec time-lapse microscopy images of HeLa cells co-expressing RAB GTPases and SNX8 and infected with *Shigella*: **A**. RAB5A-GFP/SNX8-mApple, **C**. RAB7A-GFP/SNX8-mApple, **E**. SNX8-EGFP/RAB8A-mApple, **G**. SNX8-EGFP/RAB11A-mApple. The infection foci and a zoom on a macropinosome of the infection site are shown together. Scale bars are 3µm and 1.5µm, respectively. **E**. Relative fluorescence intensity of SNX8-mApple and RAB5A-GFP to individual IAMs (n=8 events). **F**. Relative fluorescence intensity of SNX8-mApple and RAB7A- GFP to individual IAMs (n=8 events). **G**. Relative fluorescence intensity of SNX8-EGFP and RAB8A-mApple to individual IAMs (n=6 events). **H**. Relative fluorescence intensity of SNX8- EGFP and RAB11A-mApple to individual IAMs (n=8 events). In all cases, fluorescence intensity was normalized to the maximum and minimum intensities measured for each individual IAM.

RAB11A and RAB8A enrichment to IAMs was found to promote efficient BCV rupture and egress (Weiner et al. 2016, Chang et al. 2020). Therefore, we wanted to compare SNX8 recruitment with these factors. Analysis of the time-lapse images showed that in the case of RAB11A and RAB8A overexpression, SNX8 was only partially recruited to the formed IAMs (**Figures 5E** and **5G**). We distinguished in both cases 3 distinct recruitments: (i) the recruitment of both overexpressed proteins, (ii) recruitment of the RAB protein only and (iii) an exclusive SNX8 recruitment. For IAMs that showed recruitment of both host factors, we observed RAB8A recruitment to IAMs prior to SNX8 recruitment (**Figure 5E** and **5F**), however these factors did not seem to overlap at the monitored IAMs. When co-expressing RAB11A and SNX8, SNX8 was recruited first to IAMs and was then replaced by RAB11A (**Figure 5G**).

Analysis of individual IAMs also showed this switch (**Figure 5H**). Altogether, our time resolved data shows distinct profiles of SNX8 IAMs with individual Rab GTPases. Overall, these results show SNX8 recruitment is mutually exclusive to RAB11A and RAB8A at IAMs and this hints at the existence of divergent IAM subpopulations during *Shigella* infection.

### SNX8/PI(3)P impairment to IAMs hampered *Shigella* BCV egress

We proceeded to investigate the function of this newly identified subset of PI(3)P+/SNX8+ IAMs to determine their impact on BCV rupture. Since the recruitment of SNX8 was PI(3)P- driven, we assessed whether this process required the host-cell PI(3)-kinase. This was confirmed by using the broad spectrum PI(3)-kinase inhibitor wortmannin (**Supplementary Figure 1**), which led to an arrest of SNX8 recruitment to IAMs. We remarked however that SNX8 recruitment to the BCV remained, implying a PI(3)-kinase independent recruitment to the bacterial compartment.

Given the PI(3)P-dependent recruitment of SNX8 together with RAB5A, we reasoned that the RAB5 effector class III PI(3)P kinase VPS34 (Vacuolar Sorting Protein 34) was involved as it has a reported role in early endosome and macropinosome PI(3)P maturation (Bohdanowicz et al. 2013, Spangenberg et al. 2021). To test this hypothesis, we used the VPS34-specific inhibitor (SAR405) has been described to impair VPS34 function in canonical macropinosomes (Ronan et al. 2014, Spangenberg et al. 2021). Therefore, we performed time-lapse infections in HeLa cells transiently expressing mOrange-Galectin-3 and SNX8- eGFP with and without SAR405. We observed that addition of SAR405 hampered the recruitment of SNX8-eGFP to IAMs (**Figure 6A**) but did not impact bacterial invasion (**Supplementary Figure 4**). The inhibitor also impaired PI(3)P-recruitment to IAMs, but did not fully abolish the presence of PI(3)P at the BCV (**Supplementary Figure 4**).

**Figure 6.**
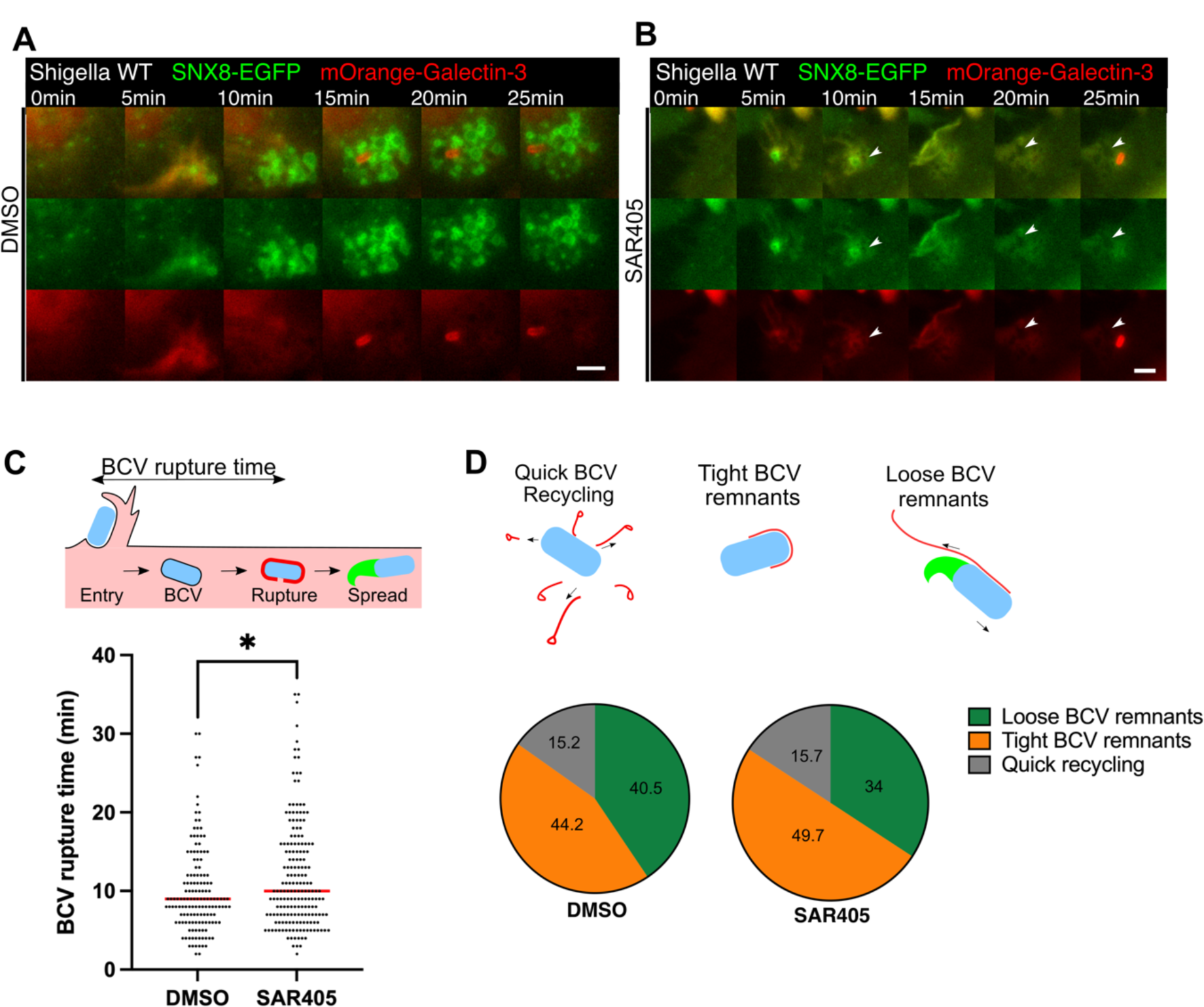
**SNX8-PI(3)P IAMs promote BCV rupture and bacteria rapid escape from the BCV. A**, B. Time-lapse images of *Shigella* infections in SNX8-eGFP/mOrange-Galectin-3 transiently transfected HeLa cells in the presence of DMSO or SAR405 inhibitor. White arrowheads show SNX8-negative IAMs. Scale bar is 5µm. **C**. BCV rupture time of the first 3 bacteria entering a bacterial focus in WT *Shigella* infected cells in the presence of DMSO or SAR405 (n>140 events for each condition). Statistical analysis was performed using an unpaired t- test (\**P*<0.05). **D**. Quantification of BCV unpeeling events in SNX8-EGFP and mOrange-Galectin-3 expressing HeLa cells treated with SAR405 or DMSO (nDMSO=233, nSAR405=289). Schematic illustrations depict the phenotypes analyzed: quick recycling with quick BCV recycling with rapid recycling of the membrane from the bacteria, tight BCV remnants with delayed bacteria movement and loose BCV remnants with bacteria movement.

Given the dynamics of SNX8 recruitment in relationship to IAM formation during *Shigella* intracellular niche formation, we examined if SNX8-IAMs function in BCV rupture or in steps of BCV disassembly. To track the BCV rupture and membrane remnants, we performed time lapse infections of SAR405 treated HeLa cells co-expressing SNX8 together with the fluorescent reporter Galectin-3. BCV rupture time was defined as the time in between membrane ruffling and Galectin-3 recruitment to the BCV (see illustration in **Figure 6C**).

Analysis of the BCV rupture time showed a slight, yet significant delay in the presence of the SAR405 inhibitor compared to the DMSO control (**Figure 6C**).

To monitor unpeeling of BCV membrane remnants, we were able to discern three phenotypes on the basis of BCV membrane movement and the swiftness of bacterial movement: we distinguish a quick BCV disassembly, loose BCV remnants leading to a rapid onset of *Shigella* motility, and tight BCV remnants with delayed intracellular *Shigella* motility (see **Figure 6D**). Quantification of these phenotypes is in agreement with previously reported proportions of each type of BCV disassembly in the presence of DMSO (Kühn et al. 2020) with tight BCV remnants: 44.2%, loose BCV remnants: 40.5% and quick recycling: 15.4%) (**Figure 6D**). We observed no change in the quick BCV disassembly phenotype in the presence of the VPS34 inhibitor (DMSO=15.2%, SAR405=15.7%,). However, the addition of the VPS34 inhibitor resulted in a shift in the efficiency of BCV disassembly (**Figure 6D**). Here, only 34% of infected cells were able to disassemble efficiently the BCV remnants, while 50% remained trapped within BCV remnants. This contrasted with untreated cells that displayed approximately equal distribution of the different intracellular localizations. Together, these results indicate a contribution of this specific subtype of IAMs (PI(3)P+/SNX8+) in promoting efficient *Shigella* BCV egress to reach the host cytosol.

## Discussion

Following a comprehensive screen of the involvement of BAR proteins during *Shigella* invasion combining time-resolved fluorescence microscopy with genetically-encoded markers for intracellular bacterial localization (Sanchez et al. 2021), we discovered the existence of distinct *Shigella*-IAM subpopulations. We studied a subset of IAMs which are PI(3)P and SNX8 positive showing their implication in efficient bacterial vacuole egress upon initial BCV breakage. We identified the PI(3)P signal to be VPS34-dependent on the IAMs (in contrast to the BCV), and we showed that at least part of this subset of IAMs underwent a RAB11A switch. Lastly, PI(3)P synthesis arrest to these IAMs shifted the balance of pathogens being trapped within broken BCVs to efficiently disassembled BCVs.

*Shigella* invasion steps require extensive membrane remodeling. Hence, monitoring BAR domain-containing factors involved in *Shigella* invasion appeared as promising. Previously, TOCA-1 was reported to be recruited to the *Shigella* actin cage and to be crucial for actin-tail formation (Leung et al. 2008, Baxt and Goldberg, 2014, Kühn et al. 2020) which we also observed (data not shown). We found an enrichment of SNX-BAR proteins, in particular SNX8 localized to PI(3)P positive IAMs and was retained until late *Shigella* invasion steps (**Figure 2**). With our data, we could determine the existence of a PI(3)P+/SNX8+ subset of *Shigella*-IAMs (**Figure 4**). Our results align with a report from Weiner et al (2016) which showed PI(3)P-labelled IAMs partially co-localizing with the fluid phase marker dextran.

Previously, macropinocytosis was proposed as the entry mechanism for bacterial pathogens (Cossart and Sansonetti, 2004). However, *Shigella*-IAMs were determined to be distinct morphologically and in composition from the bacterial phagosome-like vacuole (Weiner et al. 2016) prompting their formation as being driven by separate mechanisms. Furthermore, contact sites between the BCV and IAMs were reported (Weiner et al. 2016) as well as the recruitment of host factors RAB11A, RAB8A and the exocyst complex (Mellouk et al. 2014, Weiner et al. 2016, Chang et al. 2020) which were revealed to promote *Shigella*-BCV egress. Together these data highlight IAMs as a separate compartment with an important contribution to the infection process. Although morphologically comparable to canonical macropinosomes, the similarity in composition and formation of IAMs to “classical” macropinosomes has remained unclear. A recent study (Spangenberg et al 2021) found macropinosome maturation to be VPS34-dependent with inhibition of VPS34 leading to the refusion of macropinosomes with the plasma membrane, via RAB10 and RAB11A recruitment. Here, our results contrasted with those of Spangenberg et al with the VPS34- mediated PI(3)P+/SNX8+ IAMs subset maturing to RAB11A, highlighting how these pathogen-controlled compartments differ from canonical macropinosomes. This suggests there are additional assembly components assembled on these pathogen-controlled compartments which lead to a distinct trafficking outcome in comparison to canonical macropinosomes.

Through our study of the PI(3)P+/SNX8+ subpopulation of IAMs, we can speculate about the potential function(s) and maturation of the different subsets of IAMs. Based on RAB recruitment, we propose that some subsets may follow the RAB7A endosomal degradation pathways, SNX8+/RAB11A- subsets of IAMs may be subject to communicate with the TGN whereas SNX8-/RAB11A+ IAMs may become “recycling endosome-like” potentially undergoing recycling at the plasma membrane. This finding work highlights Shigella infection foci as being comprised of subpopulations of IAMs with potential to subvert multiple host trafficking pathways. This is in agreement with previous studies that demonstrated a function of IAMs in accelerating *Shigella*-BCV egress (Mellouk et al. 2014, Weiner et al. 2016, Chang et al. 2020) and emphasizing both a contribution of the host and the bacteria, this work showcases the *Shigella* focus as being comprised of subpopulations of IAMs evidencing the subversion by the bacterium of multiple host pathways.

Broadly our work highlights the importance of the endosomal recycling pathways in the later steps of *Shigella* invasion. *Shigella* has been shown to require rapid loss of host membrane remnants to gain access to the host cytosol (Chang et al. 2020, Kühn et al. 2020). BCV egress, has been found to require first the rupture of the BCV followed by the unpeeling of the BCV remnants. Furthermore, BCV unpeeling has been described as impacting the capacity of the bacteria to move within cells and into neighboring cells (Kühn et al. 2020, Chang et al. 2020). It is likely that the efficiency of these events impact on intracellular detection by xenophagy (Wandel et al., 2017). We show in this study, that IAMs play a role in the shedding of the BCV membrane from the bacteria. However, we show here that *Shigella* triggers maturation of different IAMs subsets with different trafficking pathways that may contribute to additional aspects of the infection process.

The bacterial factors that dictate the diversity of IAMs during *Shigella* entry need to be determined. The Δ*ipgD* mutant leads to a slight delay in bacterial entry, however it almost entirely abrogated the formation of IAMs (Garza-Mayers et al. 2015, Weiner et al. 2016). These studies together with our data using the PI(3)P kinase inhibitors suggest it unlikely that IpgD is key to controlling the different IAM subsets. Interestingly, another effector has been shown to modulate small host GTPases. IcsB is an N-fatty acylase that covalently binds small GTPases to host membranes during *Shigella* invasion (Liu et al. 2018). We have also found that it is involved in the formation of the actin cocoon, and it has an impact on egress of *Shigella* from its vacuole (Kühn et al. 2020). Therefore, it would be interesting to study how the Δ*icsB* mutant affects the different subpopulations of IAMs. Furthermore, IpaB and IpaJ have been identified to be involved in Golgi fragmentation, and it is possible that these pathways also regulate the various subsets of IAMs (Burnaevskiy et al. 2013).

Other bacterial pathogens have also been described to trigger macropinosome-like compartments during their invasion. In particular, *Salmonella* -a bacterial pathogen closely resembling *Shigella*- has been shown to form IAMs during infection. Recently, these *Salmonella*-IAMs were shown to be critical for *Salmonella* niche establishment (Stévenin et al. 2019). IAMs were shown to fuse to the *Salmonella* vacuole, forming the *Salmonella* replicative niche (90% of events), whereas impairment of IAM fusion lead to rupture of the *Salmonella* vacuole (Perrin et al. 2004, Malik-Kale et al. 2012, Knodler et al. 2014). Hence, a comparison of the composition of *Salmonella*-IAMs and *Shigella*-IAMs is crucial to understanding bacteria niche establishment. This could help to highlight specific pathways exploited by bacterial pathogens to establish their intracellular niche.

SNX-BAR proteins have been previously shown to be important targets for invasive bacteria. The *Chlamydia* “inclusion”, the pathogen’s replicative niche, was reported to selectively recruit SNX5, SNX6 and SNX32 through the *Chlamydia* effector IncE. These were shown to form tubules at this compartment and to be crucial for Chlamydia replication within its host (Mirrashidi et al. 2015, Elwell et al. 2017, Paul et al. 2017). *Salmonella,* a bacterium residing within a replicative niche called the *Salmonella*-containing vacuole (SCV), has also been reported to reprogram SNX-BAR proteins. *Salmonella* has been shown to hijack SNX-18 for its entry within the host cell (Liebl et al. 2017) and once internalized, *Salmonella* induces SNX recruitment to the SCV through the *Salmonella* effector SopB-mediated by PI(3)P formation (Stévenin et al. 2019). Similar to *Chlamydia*, SNX1 forms tubules at the SCV which have been shown to regulate bacterial niche establishment (vacuole stability or rupture).

SNX-BAR subversion has also been reported in a cytosol-residing bacterium. Through the oral listeriosis enhancing effector Lmo1656, *Listeria* has been described to recruit SNX6 for successful *Listeria* infection (David et al. 2018). Altogether these discoveries highlight that SNX-BAR proteins have an important role for membrane remodeling during bacterial invasion across multiple species. From this it is clear that further studies to define how these proteins influence pathogenesis at the host-microbe interface are needed, as they may represent a potentially novel target to prevent bacterial niche establishment.

## Authors contribution

Conceived and designed all experiments: LS, CV and JE. Performed and analyzed data for all experiments: LS with specific contributions from MGC (pSBbi-SNX8-EGFP and pSBi-LactC2- EGFP cloning and in SNX8-EGFP cell line development). LS, CV and JE wrote the original draft of the manuscript, and LS, CV, MGC, MH and JE edited this manuscript. CV and JE supervised the research and JE secured funding.

## Acknowledgments

We would also like to thank Magdalena Gil for feedback on this manuscript and Yuen-Yan Chang, Laura Barrio-Cano and Sonja Kühn for fruitful discussion on this project. We would like to thank Laurent Audry for technical support. We thank Sandrine Schmutz of (UTechS Cytometry and Biomarkers) of C2RT for technical assistance during cell sorting.

## Supplementary data

**Supplementary Figure 1.**
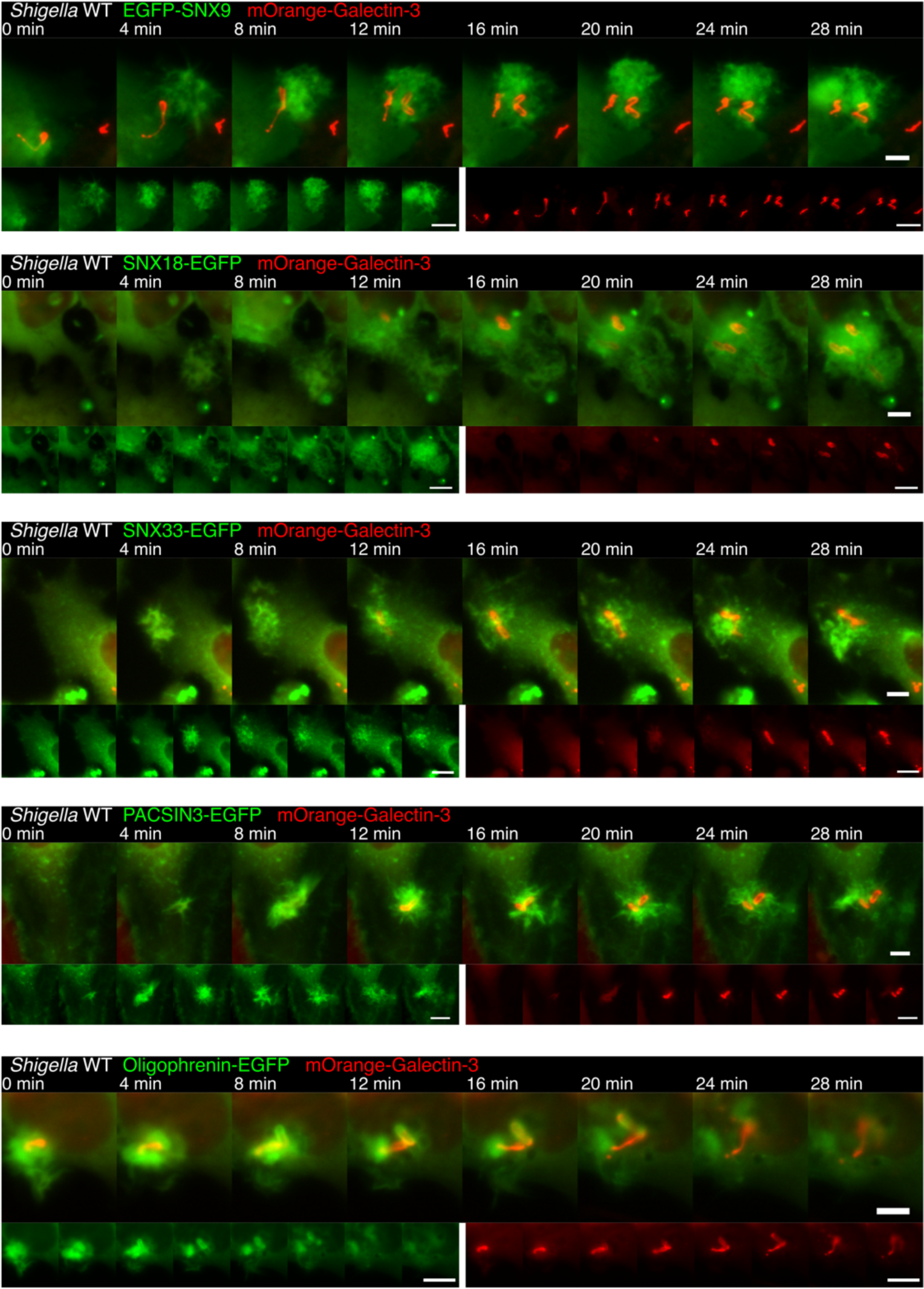
Example of BAR domain protein hits during screening. HeLa mOrange-Galectin-3 transfected with either EGFP-SNX9, SNX18-EGFP, SNX33-EGFP, PACSIN3-EGFP or Oligophrenin-EGFP were infected with *Shigella* WT. mOrange-Galectin-3 was used a reference point and marks the breakage of the *Shigella*-BCV. Scale bar is 5µm and 10µm.

**Supplementary Figure 2.**
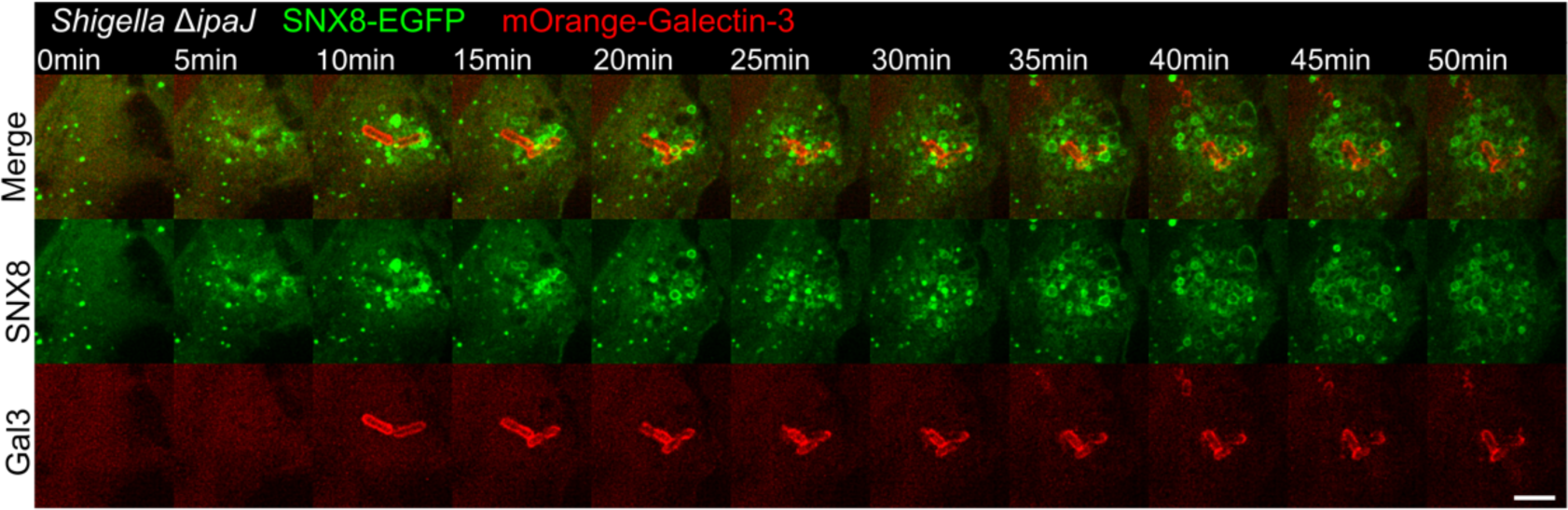
*Shigella ΔipaJ* infection control. HeLa cells expressing mOrange-Galectin-3 and SNX8-EGFP were infected with *Shigella ΔipaJ*. mOrange-Galectin-3 marks the breakage of the *Shigella*-BCV. Scale bar is 5µm.

**Supplementary Figure 3.**
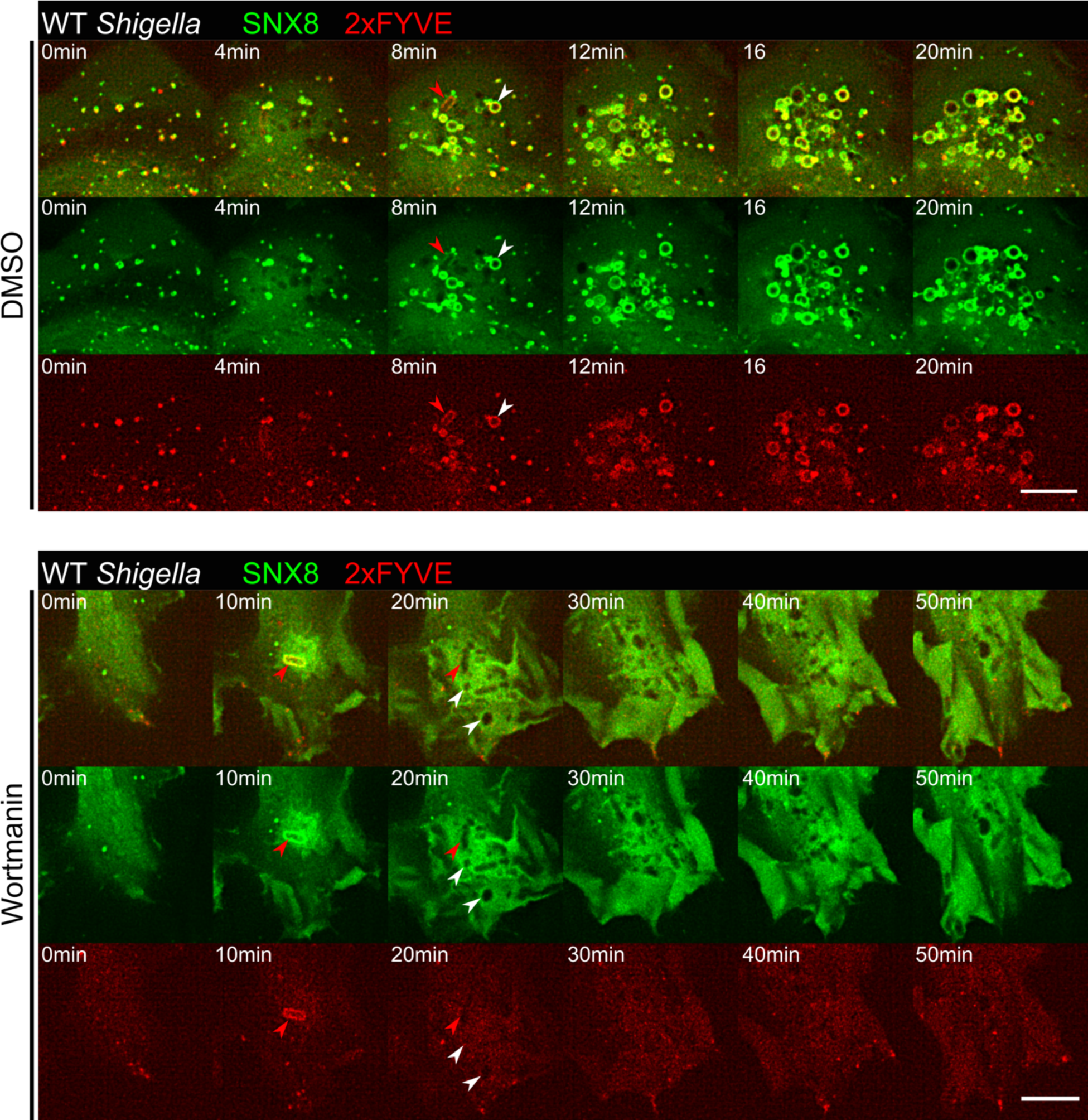
Wortmannin inhibition of PI(3)P *de novo* synthesis leads to arrest of SNX8 recruitment to IAMs HeLa mOrange-Galectin-3 were transfected with SNX8 and infected with wildtype *Shigella* together with wortmannin (0.44µM). Red arrowheads show an example of entering bacteria. White arrowheads show examples of formed IAMs. Scale bar is 5µm.

**Supplementary Figure 4:**
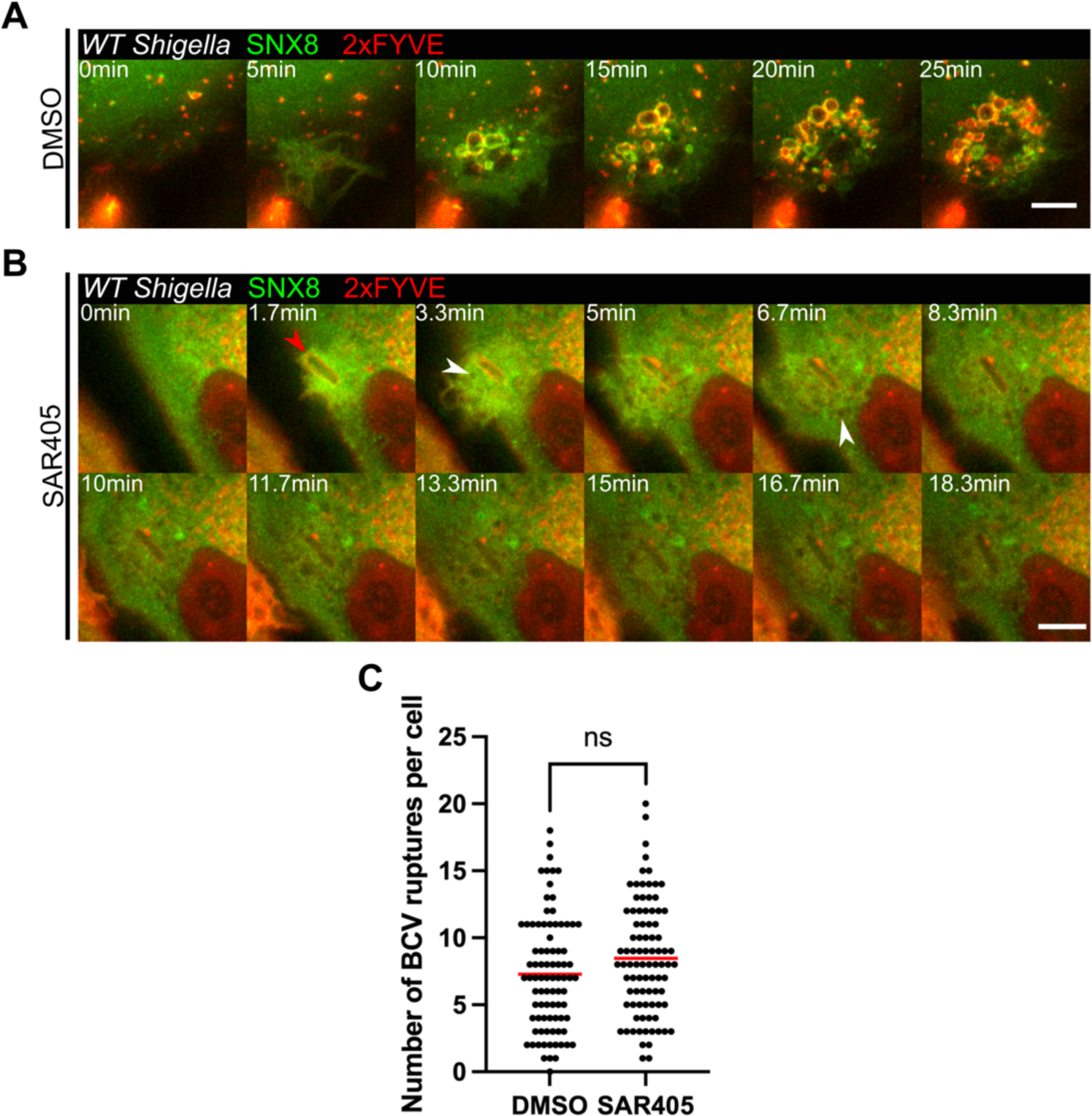
Effect of SAR405 on PI(3)P presence at the *Shigella* infection focus. A, B. Microscopy images of HeLa cells co-transfected with SNX8-eGFP and 2xFYVE-mCherry infected with WT *Shigella* treated with DMSO or PI(3)-kinase VPS34 inhibitor SAR405. Red and white arrows show an entering the bacterium and a formed macropinosome, respectively. Scale bar is 5µm. C. Quantification of the number of BCV ruptures observed per cell in the presence or absence of SAR405. The quantification was performed on 3 independent experiments (n>80). A *t*-test was performed to determine the statistical significance of the data (ns *P*>0.05).

**Supplementary Table 1.**
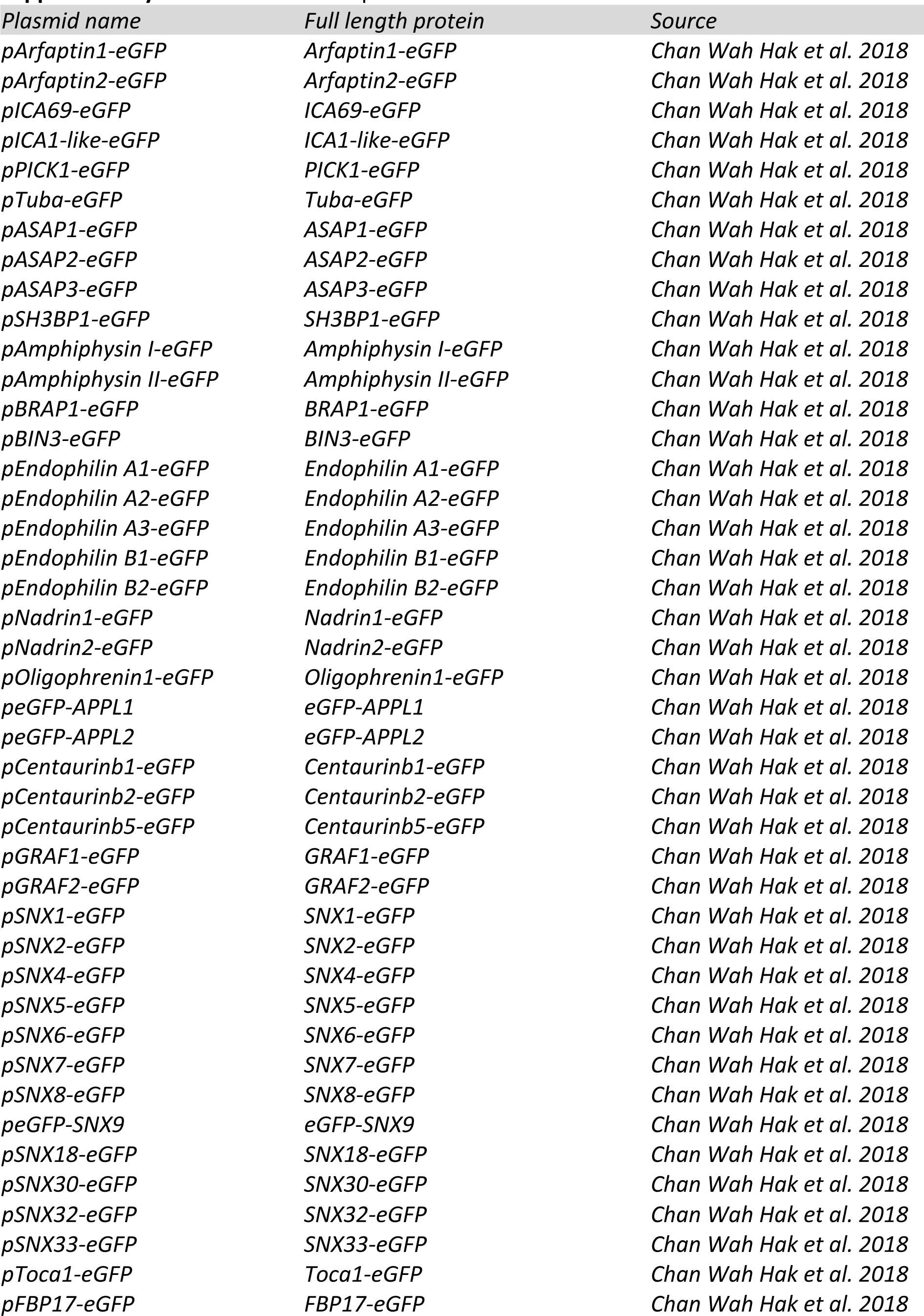
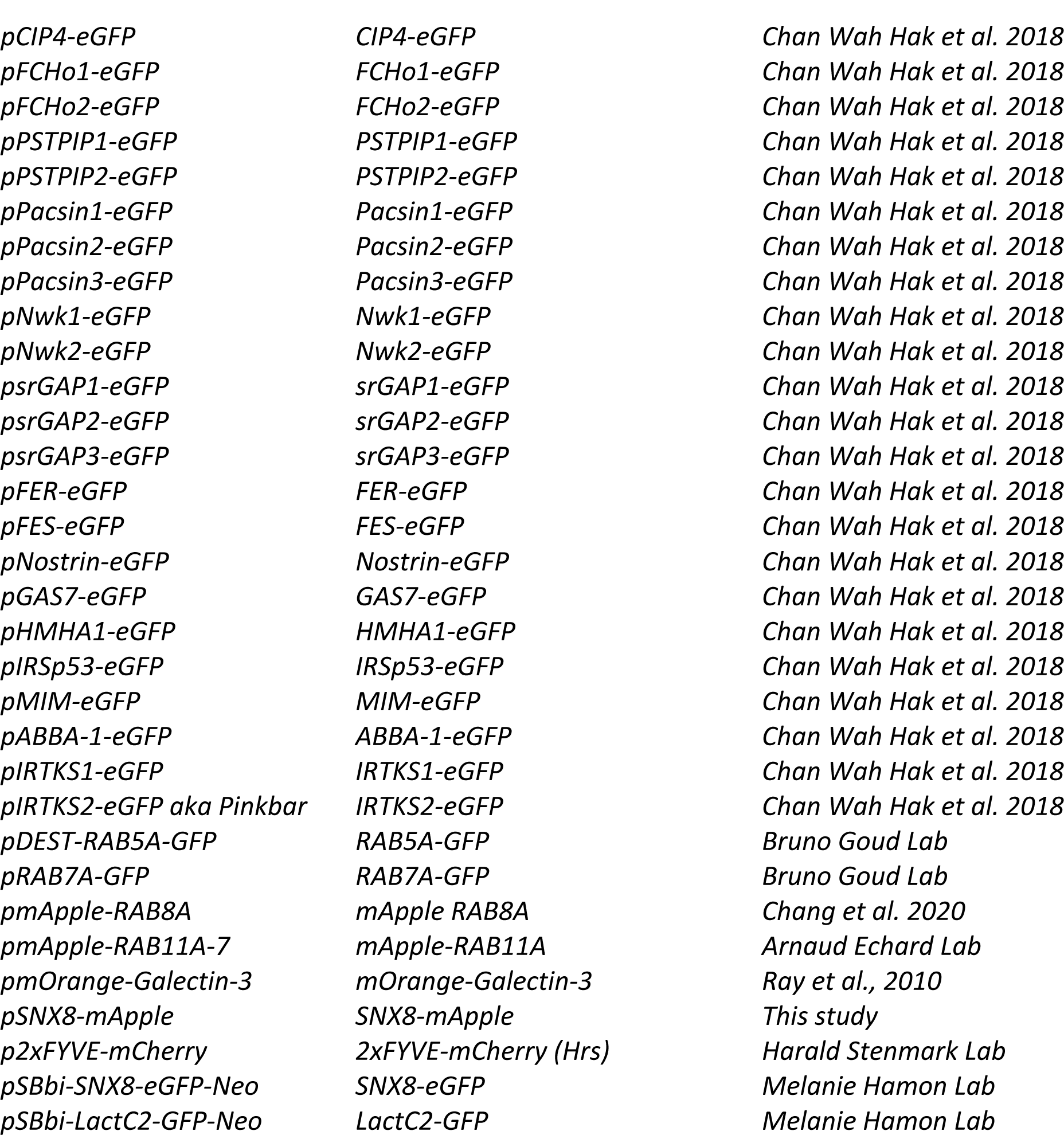
List of DNA plasmid constructs used.

**Supplementary Table 2.**
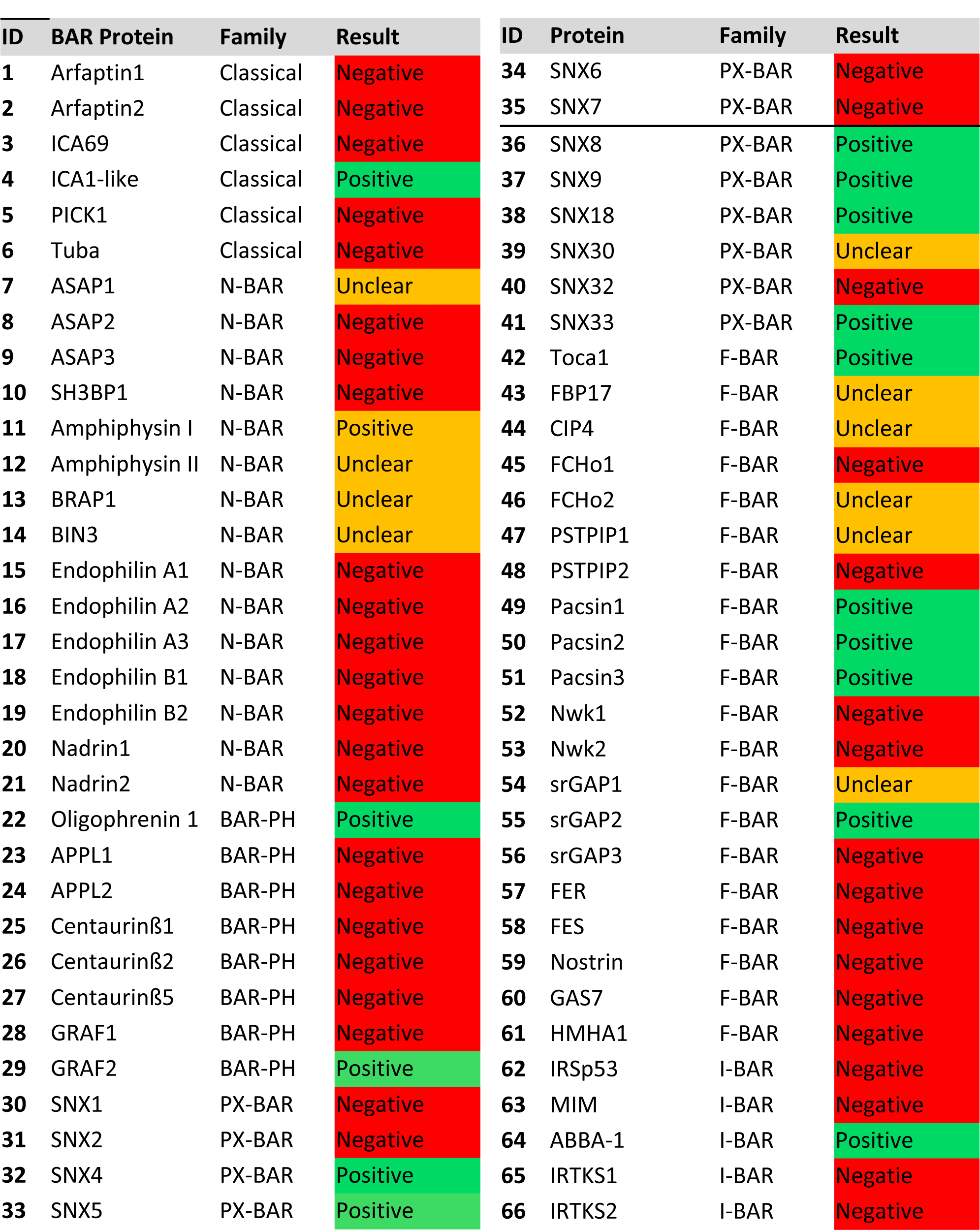
**Summary of the BAR domain-containing protein screen results.** The results shown are a summary of 3 independent experiments on each candidate protein. Positive hits were counted as being observed to the infection site at least twice, negatives being all 3 tries showed no visible recruitment and unclear being that recruitment was observed only once.

